# Optimal motion-in-depth estimation with natural stimuli

**DOI:** 10.1101/2024.03.14.585059

**Authors:** Daniel Herrera-Esposito, Johannes Burge

## Abstract

Estimating the motion of objects in depth is important for behavior, and is strongly supported by binocular visual cues. To understand both how the brain should estimate motion in depth and how natural constraints shape and limit performance, we develop image-computable ideal observer models from naturalistic binocular video clips of two 3D motion tasks. The observers spatio-temporally filter the videos, and non-linearly decode 3D motion from the filter responses. The optimal filters and decoder are dictated by the task-relevant natural image statistics, and are specific to each task. Multiple findings emerge. First, two distinct filter types are spontaneously learned for each task. For 3D speed estimation, filters emerge for processing either changing disparities over time (CDOT) or interocular velocity differences (IOVD), cues used by humans. For 3D direction estimation, filters emerge for discriminating either left-right or towards-away motion. Second, the filter responses, conditioned on the latent variable, are well-described as jointly Gaussian, and the covariance of the filter responses carries the information about the task-relevant latent variable. Quadratic combination is thus necessary for optimal decoding, which can be implemented by biologically plausible neural computations. Finally, the ideal observer yields non-obvious–and in some cases counter-intuitive–patterns of performance like those exhibited by humans. Important characteristics of human 3D motion processing and estimation may therefore result from optimal information processing in the early visual system.

**Significance statement:** Humans and other animals extract and process features of natural images that are useful for estimating motion-in-depth, an ability that is crucial for successful interaction with the environment. But the enormous diversity of natural visual inputs that are consistent with a given 3D motion–natural stimulus variability–presents a challenging computational problem. The neural populations that support the estimation of motion-in-depth are under active investigation. Here, we study how to optimally estimate local 3D motion with naturalistic stimulus variability. We show that the optimal computations are biologically plausible, and that they reproduce sometimes counterintuitive performance patterns independently reported in the human psychophysical literature. Novel testable hypotheses for future neurophysiological and psychophysical research are discussed.

## Introduction

Motion in depth, relative to an observer, can be generated by self-motion or by the motion of objects in the world. Accurate estimation of motion in depth is important for successful interaction with the environment. Animals with binocular vision combine the two two-dimensional (2D) images formed by the eyes to estimate 3D motion. However, many questions remain about how the brain does, and should, estimate 3D motion from binocular images (Bonnen et al., 2020; Cormack et al., 2017; Czuba et al., 2014; Rosenberg et al., 2023; Sanada & DeAngelis, 2014). Ideal observer analysis is a useful tool for providing normative accounts of how perceptual tasks should be solved, and for establishing theoretical bounds on sensory-perceptual performance. Ideal observer analysis can thus aid experimentalists in developing principled hypotheses about real biological systems that can be tested empirically (Burge, 2020; Geisler, 1989).

Previous studies have applied Bayesian-observer analysis to tasks of 3D motion estimation (Lages, 2006; Rokers et al., 2018; Welchman et al., 2008). However, previous attempts to model 3D motion estimation do not account for natural (nuisance) image variability, and are not image-computable. Existing models therefore make assumptions about the properties of stimulus encoding, and the quality of the information it provides about 3D motion. These assumptions may be inconsistent with the constraints imposed by natural images and the front-end of the visual system. As a consequence, these models say little about what early visual computations–including what stimulus features–are most useful for estimating 3D motion from images, and how these computations depend on the properties of natural signals. These concerns can be alleviated by developing Bayesian-observer models that are grounded in the statistical properties of natural images and scenes.

To better understand the computations that optimize local 3D motion estimation, we developed two image-computable Bayesian ideal observer models that were constrained by the front-end of the human visual system and incorporated to the statistics of naturalistic binocular video clips. Each ideal observer was designed to perform one of two specific tasks: 3D speed estimation or 3D direction estimation. For each task, the observer model first applies a small set of task-optimized spatio-temporal filters to the binocular videos. The filters are learned via Accuracy Maximization Analysis (AMA), a Bayesian method for finding receptive fields that select for the stimulus features that provide the most useful information for a given task (Burge & Jaini, 2017; Geisler et al., 2009). Here, we use AMA-Gauss (Jaini & Burge, 2017), a computationally efficient form of AMA that makes the strong assumption the the receptive-field-responses are Gaussian distributed; this assumption is empirically justified for the 3D motion tasks being investigated here (see Results). After the task-optimized filters have been learned, the ideal observer performs optimal non-linear decoding, using the statistics of the filter responses and the tools of probabilistic inference, to yield optimal estimates of the task-relevant latent variable (i.e. 3D speed or 3D direction; **Figure 1AB**).

**Figure 1.**
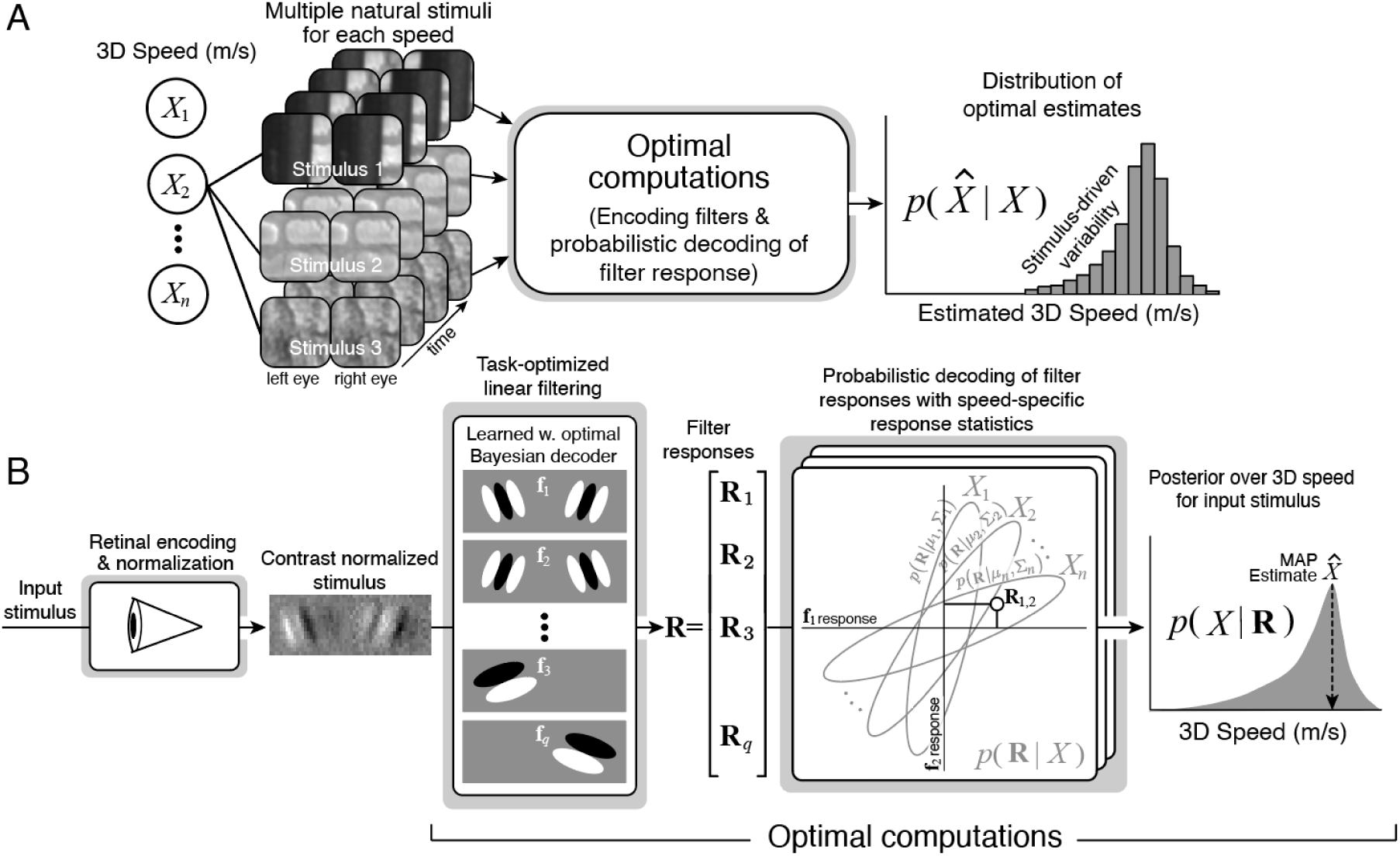
Ideal observer model for 3D speed estimation with natural stimuli. **A,** In natural viewing, many different retinal stimuli are associated with the same value of a latent variable (*X*). The naturalistic dataset used here mimics this natural (“nuisance”) variability: many different binocular image videos correspond to the same 3D motion trajectory in depth. For each binocular image video having a certain 3D motion (here, 3D speed), the ideal observer outputs an optimal estimate (*X̂*) of that speed. Across all binocular image videos having that same 3D speed, a corresponding distribution of optimal estimates results. The variance of the estimates is largely stimulus-driven, and attributable to natural image variability. **B,** Information flow through the ideal observer. First, an input stimulus is encoded by the retina and normalized to yield a contrast-normalized retinal stimulus. Then, task-optimized receptive fields are applied to the stimulus, yielding a set of filter responses (**R**). Next, each set of filter responses is decoded using the statistics–µ*_m_* and Σ*_m_*–that characterize the conditional probability distributions of response for all speeds. Finally, the resulting posterior probability distribution over 3D speed is then used to obtain an optimal estimate.

The most important results are as follows. For each task, two filter types with distinct functional specializations are learned, reflecting a spontaneously emergent division of labor. The response statistics of the learned filters to naturalistic videos dictate that optimal 3D speed and 3D direction estimation require computations that can be well-approximated by an extension of the widely studied energy model of cortical neural responses (Adelson & Bergen, 1985; DeAngelis et al., 1991; Ohzawa et al., 1990). Finally, for both 3D speed and direction estimation, we compare the performance of the two image-computable ideal observer models against previously reported human psychophysical data (Bonnen et al., 2020; Czuba et al., 2010a; Fulvio et al., 2015; Rokers et al., 2018), and show that the models provide a normative account of many aspects of human psychophysical performance.

## Methods

### 3D motion estimation tasks

We trained observer models on two different 3D-motion estimation tasks with naturalistic binocular videos: 3D-speed estimation and 3D-direction estimation. To train each observer, the model is presented with a labeled set of input binocular videos, each consistent with a different rigid planar surface moving in 3D space, behind a fixed circular aperture, with a given speed and direction (**Figure 2A**). The objective was to estimate, from the input videos, the 3D motion of the surface. A dataset of binocular video clips with known 3D motion was generated for each task.

**Figure 2.**
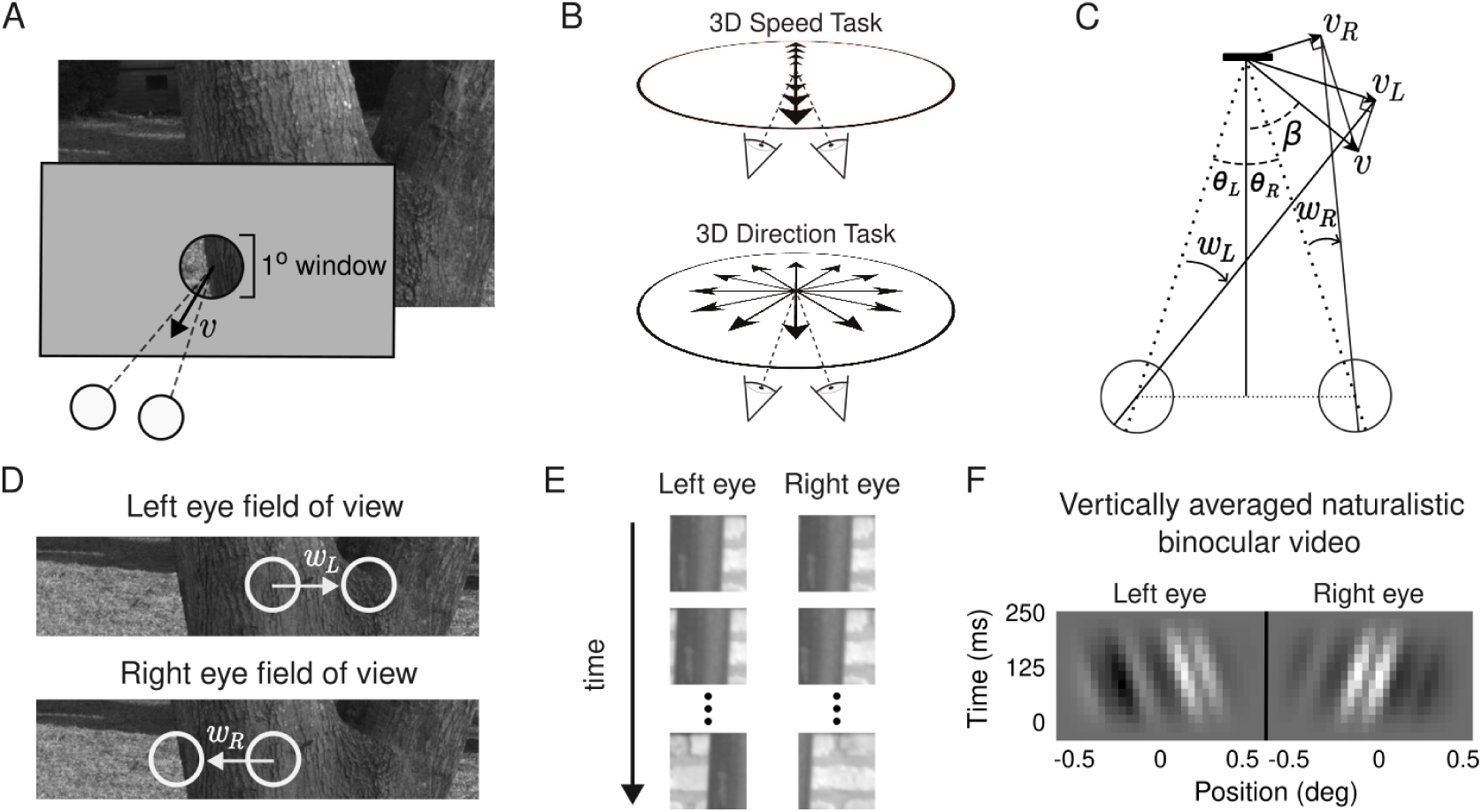
Task design and stimulus synthesis. **A,** Visualization of the information available to the ideal observer. A flat, rigid, surface moves in 3D as indicated by the solid motion vector *ν*. The observer views the surface from behind a fixed 1 degree window. **B,** (Top) Observer looking at the target surface (black bar) for the speed estimation task. Arrows show the types of 3D motions used in this task. (Bottom) Same but for the direction estimation task. **C,** Projective geometry linking 3D motion (*ν*) to retinal speeds (ω_*L*_, ω_*R*_) in degrees of visual angle per second. **D,** Videos are synthesized by translating the field of view of each eye in the target surface according to the corresponding retinal speeds ω*_L_* and ω_*R*_. The region of the target surface visible to each eye changes as the surface moves in 3D. **E,** Example of naturalistic binocular video. The first frames of the left eye and right eye videos are identical because the initial disparity is zero (top), but image-features drift apart with time (bottom). Note that after sufficient time has elapsed, there are few if any corresponding points in the left- and right-eye images. **F,** Example of vertically averaged naturalistic binocular video, moving towards the observer at 1.4 m/s. Each horizontal slice of the stimulus corresponds to a frame of a binocular video (see E) that was vertically averaged. The stimulus includes pixel noise and is contrast normalized.

In the 3D-speed estimation task, each video shows a planar surface starting from 1 m in front of the observer and moving directly towards or away from the observer with a given speed (**Figure 2B top**). The set of speeds used in the videos range from -2.5 m/s (receding) to 2.5 m/s (approaching) in 0.1 m/s increments (51 total speeds). These 3D speeds correspond to monocular retinal speeds ranging from -4.6 to 4.6 deg/sec. Thus, the task for the model is inferring which of the 51 possible speeds is shown in the binocular video. In the 3D-direction estimation task, each video shows a surface starting 1 m straight ahead of the observer, and at a fixed speed of 0.15 m*/*s in some direction in the XZ plane (**Figure 2B bottom**). Directions covering the whole XZ plane in 7.5 deg increments were used (48 total directions), and the task consists in inferring the direction of the target surface.

### Binocular video synthesis

We generated a large number of naturalistic binocular videos for each 3D motion. The variation across videos constitutes naturalistic (‘nuisance’) stimulus variability to the dataset and the task. For each naturalistic video we first sampled a different patch from a dataset of natural scenes (Burge et al., 2016) to present as the moving surface (**Figure 2AD**). For each 3D motion, we computed the corresponding retinal velocities in the left- and right-eye images using standard projective geometry. The left- and right-eye velocities are given by

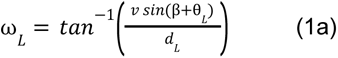

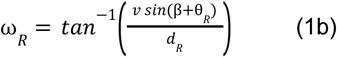

where ω_*L*_ and ω_R_ are, respectively, the left- and right-eye angular velocities in degrees per second, *ν* is the speed of the target, □ is the target direction with respect to the midsagittal plane, *θ*_L_ is the angle between the left-eye and cyclopean line-of-sight, and *θ*_R_ is the analogous angle for the right eye (**Figure 2C**). Then, for each eye, we generated a video by taking crops of the target patch that were translated at each frame in accordance to the retinal speed (**Figure 2D**). The videos for the two eyes start from the same location in the natural scene, and thus the binocular video starts with zero disparity (**Figure 2E**, first panel in each row). When the displacement between frames included fractions of a pixel, pixels were interpolated accordingly. We also incorporated the effects of the well-focused human eye (Thibos et al., 1992; Wyszecki & Stiles, 1982), and the spatial sampling, photopic (L+M) sensitivity, and temporal impulse–response function of the cone photoreceptors (Curcio & Allen, 1990; Schneeweis & Schnapf, 1995; Stockman & Sharpe, 2000). (See Burge & Geisler, 2011, 2014, 2015 for more detailed characterizations of these procedures.) Videos were then downsampled to 30 pixels/deg. (We have previously verified that, because the most useful information for the task resides in lower spatial frequencies, downsampling has little practical effect on performance.) Binocular videos had a duration of 250 ms (60 Hz) and subtended 1 deg of visual angle in each eye (30 pix/deg). This field of view, which was enforced by a raised cosine spatio-temporal window, approximates the information received by a receptive field in the early visual cortex.

Each frame of the video was then vertically averaged to produce a 1D binocular video (**Figure 2F**). This analytic choice was made because it vastly reduces the computational cost of the filter learning procedure. Vertical averaging the stimuli does not change the response of vertically-oriented filters because such filters themselves vertically average the stimuli (see Burge & Geisler, 2015). Indeed, it is the computational equivalent to restricting the filter learning procedure to return only vertically oriented receptive fields. The ideal observers described in the article can be conceived of as operating on the outputs of receptive fields within vertically-oriented orientation columns. A clear direction for future work is to generalize the analysis to videos that are not vertically averaged. Technical improvements in the computational efficiency of receptive-field learning will facilitate these efforts (see Burge & Jaini, 2017; Jaini & Burge, 2017). Such improvements are well underway.

The binocular intensity videos were then converted to binocular contrast videos by subtracting and dividing by the mean intensity of each video. A given binocular video obtained with this process (**Figure 2F**) is represented (for training or testing) as a vector with 900 elements (2 eyes x 30 pixels x 15 frames). For both tasks we generated 800 binocular videos for each motion (**Figure 1A**), 500 for training and 300 for testing the model.

We supplemented the main stimulus sets with additional stimulus sets that included looming cues, and that included disparity variability. Disparity variability was introduced by including random variability in the initial disparity of the videos (instead of starting always from zero disparity). In each such binocular video we sampled an initial binocular disparity from a normal distribution with zero mean and a given standard deviation. This initial disparity was achieved by starting the videos of each eye from locations offset in depth from fixation, with the offset corresponding to the initial disparity. Second, we generated datasets such that included looming cues, which are the changes in retinally-projected size that occur as objects move towards or away the observer. To do so, we computed, using projective geometry, the pixel-by-pixel pattern of retinal image motion that would be caused by a plane moving through 3D space. With this procedure, images of features on the target surface subtend larger or smaller visual angles (i.e. portions of the visual field) depending on the instantaneous position of the surface, relative to the observer, in 3D space.

### Ideal observer model

The stimulus synthesis procedure above generates a set of binocular videos, where each video shows a surface moving with a given 3D motion. We refer to the latent variable that is the 3D motion in each video as *X*, which can take values of *X*_1_, … , *X_n_*. For example, in the 3D speed estimation task, *X* is the speed of the surface, and *X*_*i*_ is a given speed value. As described above, for each motion *X*_*i*_ in a task we synthesized a set of binocular videos, ***s***_*ij*_, where *i* is the index of the true 3D motion and *j* indexes the different videos corresponding to motion *i* (see **Figure 1A**). To solve the task of assigning the correct label *X*_*i*_ to stimulus ***s***_*ij*_, we train an ideal observer model using Accuracy Maximization Analysis (AMA), which has been used to train similar models for related visual tasks. The ideal observer model has three different stages: pre-processing, encoding and decoding (see **Figure 1B**).

The pre-processing stage involves biologically realistic steps similar to those occurring in the retina and early visual cortex. First we add a sample of white noise γ ∼ *N*(0, σ_*p*_) to each pixel. Then, consistent with standard models of neural response (Albrecht & Geisler, 1991; Burg et al., 2021; Heeger, 1992; Iyer & Burge, 2019), the stimulus is contrast-normalized as follows:

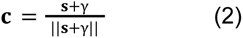

Normalization is done separately for each eye.

During the encoding stage, a small set of linear spatiotemporal filters is applied to the noisy normalized stimuli *c,* and a sample of Gaussian noise is added to the response of each filter to obtain a scalar noisy filter response. Each filter is constrained to have unit magnitude. For an individual filter **f**_*l*_, its noisy response is given by.

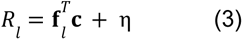

where η ∼ *N*(0, σ_r_^2^) is a sample of Gaussian noise. Denoting the set of filters **f** = [**f**_1_, … , **f**_*q*_] and population filter response vector **R** = [*R*_1_, … , *R*_q_] ∈ ℝ*^q^* , the noisy filter responses, conditional on a particular stimulus 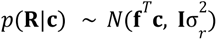, are normally distributed with filter noise having a diagonal covariance matrix Iσ_r_^2^. The noisy responses of the linear filters can be conceptualized as the normalized drives to simple cells in the visual cortex. However, the stimulus-conditioned response distributions *p*(**R**|**c**) are not particularly useful in isolation for estimating 3D motion, because the same 3D motion can cause an infinite number of natural binocular videos. Much more useful are the 3D-motion-conditioned response distributions 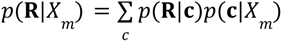 which can be obtained by marginalizing out all the binocular videos in the dataset corresponding to a given value of the latent variable.

During the decoding stage, the value of the latent variable *X* is inferred from the vector of filter responses *R* that a stimulus caused (see **Figure 1B**). The posterior probability *p*(*X*_m_|*R*) of a latent-variable value *X_m_* given a set of filter responses can be computed as the product of the likelihood *L*(*X*_*m*_; **R**) = *p*(**R**|*X*_*m*_) and the prior *p*(*X*_*m*_). To compute the likelihood that a stimulus having arbitrary value *X*_*m*_ elicited the response, when each conditional response distribution is Gaussian-distributed, *p*(**R**|*X*_*m*_) ∼ *N*(µ_*m*_, Σ_*m*_), the value **R** is plugged into the equation for each appropriate Gaussian. Specifically, the likelihood is given by

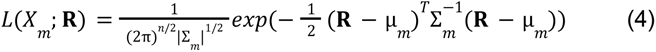

where the conditional mean *µ_m_* and covariance Σ*_m_* are estimated from the responses to all stimuli in the training set corresponding to latent variable value *X*_*m*_. Note that the log-likelihood is quadratic function in **R**, which is why energy-model-like computations are required for optimal decoding (see Results). To obtain an optimal estimate *X̂* for a given cost function, the posterior probability for each value of the latent variable *p*(*X* = *X*_*m*_|**R**) must be obtained. From the posterior probability distribution *p*(*X*|*R*), we obtain the maximum a posteriori (MAP) estimate of the value of the latent variable (i.e. the level of the latent variable with the highest posterior probability).

The standard deviation of the pixel noise in Weber contrast units was σ_*p*_ = 0. 008 and the standard deviation of the output noise associated with each receptive field was σ_r_ = 0.1. These values are in line with the signal-to-noise ratio and dynamic range of real neurons in cortex (Geisler, Najemnik, Ing, 2009). Results are also robust to the specific values chosen for these parameters over a wide range (see Burge & Jaini, 2017).

### Filter learning

The optimal filters (receptive fields) were learned using AMA-Gauss, a version of AMA (Jaini & Burge, 2017). AMA-Gauss learns filters given the (verifiable) assumption that, when conditioned on each value of the latent variable, the filter responses are Gaussian distributed. Previous work on related tasks–binocular disparity estimation and 2D motion estimation–shows that the filters learned using the assumption that the conditional response distributions are Gaussian are near-identical to those learned without the assumption (Burge & Jaini, 2017; Jaini & Burge, 2017). The loss function was chosen to be the Kullback-Liebler divergence between the computed posterior probability distribution and an idealized posterior probability distribution with all its mass at the correct value of the latent variable. In this case, the cost associated with each stimulus is given by the negative logarithm of the posterior probability at the correct value of the latent variable

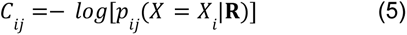

where *R* are the stochastic filter responses to the *ij*^th^ stimulus, with latent variable value of *X* = *X*_i_ .(Note that in the current case, the KL divergence is equivalent to the cross-entropy loss.) The total cost is the stimulus-specific cost averaged across all stimuli in the dataset

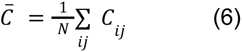

Stochastic gradient descent was used during optimization, with batches of 2048 stimuli, out of a total of 25500 and 24000 stimuli for the 3D speed and 3D direction tasks respectively. The step-size decreased by 20% every ten epochs. One hundred epochs were used. The loss always plateaued. Given the assumption that the conditional response distributions are Gaussian, the sufficient response statistics–that is, the mean and covariance *µ_j_* and Σ*_j_* of the filter responses –were re-computed after each update of the filters. The whole training set was used to update the statistics at each step.

Filters were learned in rank-ordered pairs. For each new learned pair, previously learned filters were kept fixed. We learned ten filters for each task because learning additional filters returned copies of previously learned filters, and because performance began to asymptote. (Note that when AMA returns copies of previously learned filters it signifies that the reduction of internal noise is more beneficial than the extraction of new stimulus features.)

To reduce the chance of local minima, each pair of filters was trained seven times with different starting seeds; the pair of filters with the lowest final loss was used while the rest were discarded. This procedure resulted in highly consistent sets of filters for different starting conditions.

### Interpolating response statistics

To obtain MAP estimates that were not limited by the spacing of the latent variable in the training set, we interpolated between the values used for training. Specifically, we fitted a cubic spline to each element of the sample mean and sample covariance functions of the filter responses (µ(*X*) and Σ(*X*), respectively). Then, using the cubic spline fit, we interpolated these sufficient response statistics for values of the latent variable that were not in the dataset. This procedure works well because the response statistics change smoothly with *X*, and because the values of *X* were already rather finely spaced in the training dataset. We verified the accuracy of this interpolation procedure by comparing interpolated statistics to left-out empirical statistics. We interpolated ten points in between each neighboring pair of latent variables *X*_*m*_ and _*Xm*+1_ in the training set. In the set of interpolated means and covariances, the speed increments were 0.008 m/s and the direction increments 0.625 deg for the estimation results shown. This procedure ensured that the posterior probability distributions over the latent variable, and hence the MAP estimates, were not quantized by the latent variable spacing in the training set. Results are qualitatively the same without using interpolation, although some quantization effects are apparent.

### Caveats

There are two important caveats to note before proceeding further. First, the binocular videos described above are compatible with many different 3D motions, depending on the starting location of the target (Lages & Heron, 2010; Longuet-Higgins, 1984). Information about 3D location must be available to obtain unambiguous estimates of 3D motion from such binocular videos. Just as in stereo-depth perception solutions to the stereo-correspondence problem do not address how 3D location is estimated, the current computations do not address how 3D location is estimated; we assume such information is known to the observer. However, the resulting estimates of 3D motion from the ideal observer trained on videos from one distance (1m away, straight-ahead), can be adapted for a different 3D location by appropriate geometrical transformation (see Discussion for more).

The second important caveat is that, given the design of the current analyses, the 3D speed and 3D direction tasks both could, in principle, be performed by analyzing 2D speed from one eye only. This same issue has been faced by human psychophysical studies with closely related designs. In such studies, control experiments have examined whether humans can, when tasked to do so, reliably report 2D motion in one or the other eye. Such studies have come to the justified conclusion that human 3D motion estimation and discrimination performance is not accounted for by explicit reliance on 2D motion signals in each eye alone. Stereomotion suppression (Brooks & Stone, 2006; Tyler, 1971), a general difficulty with utrocular (i.e. eye of origin) discrimination and identification (Ono & Barbeito, 1985), and various other controls contribute to this consensus (Czuba et al., 2010a; Harris & Watamaniuk, 1995; Rokers et al., 2008). The fact that humans cannot explicitly report 2D motion of course does not mean that the image-computable ideal observers developed here do not make exclusive use of 2D motion signals in each eye. But there are at least two reasons to think otherwise, and to instead conclude that binocular comparisons carry distinct information for the task. First, for each respective task, an optimization procedure that finds the most useful stimulus features returns binocular receptive fields. Second, binocular comparisons of monocular filter responses clearly carry information over and above that provided by the responses of each monocular filter in isolation. Hence, binocular comparisons substantively underlie ideal observer performance in both the 3D speed and 3D direction tasks. Nevertheless, the development of expanded observer models that jointly estimate 3D speed and 3D direction is important future work, as they will enable yet stronger conclusions to be drawn regarding the binocular computations that should underlie 3D motion estimation (see Discussion for more).

### Code

Stimuli were generated using a database of natural images (Burge et al., 2016), and custom written software in Matlab. Filter learning routines and ideal observer analyses were implemented in Pytorch. The code is available at https://github.com/dherrera1911/3D_motion_ideal_observer.

## Results

We developed image-computable ideal observer models for two different 3D motion estimation tasks with naturalistic stimuli: 3D speed estimation (**Figure 2B top**) and 3D direction estimation (**Figure 2B bottom**). The training and test dataset of naturalistic stimuli were obtained by applying the laws of projective geometry to samples from a database of images of natural scenes. The resulting binocular videos approximated the retinal stimulation that would be caused by rigid surfaces moving in depth relative to the observer. The surfaces were textured with natural scenes (**Figure 2E**), adding naturalistic nuisance variability to the task (**Figure 2A**). The stimulus set and task design included a number of simplifications that limit the naturalness of the problem considered here. Although these simplifications are broadly aligned with design elements that are common to the psychophysical and neuroscience literature on 3D motion perception, in the Discussion we examine how they limit the scope of the conclusions that can be drawn from the current results. The model’s front-end processing of the stimuli is constrained by the front-end processing of the human visual system (optics, photoreceptor response properties, contrast normalization; see Methods). The ideal observers are defined by a set of task-optimized encoding receptive fields and a probabilistic decoding stage that yield optimal 3D motion estimates for natural scenes. We analyze the computations that support ideal performance with natural stimuli, and compare ideal to previously reported human performance. The current analysis takes important steps towards a better understanding of the computations that support optimal 3D speed and 3D direction estimation in natural scenes, and of the patterns that characterize human estimation and discrimination of 3D motion.

### 3D speed estimation

We developed an ideal observer for the estimation of 3D-speed from naturalistic binocular videos. The task-optimized filters, the filter responses, and ideal observer performance are shown in **Figure 3**. The filters are Gabor-like and localized in spatio-temporal frequency, like typical receptive fields in the early visual cortex (**Figure 3A**). The filter responses, conditional on different 3D speeds, are shown in **Figure 3BC**. Responses to stimuli of the same 3D speed are approximately Gaussian-distributed. The 3D speed estimates, which are decoded by the ideal observer from the filter responses, are accurate and precise (**Figure 3D**). Also, the estimation error increases systematically with speed (**Figure 3E**), similar to how human 2D speed discrimination thresholds increase as a function of speed (Chin & Burge, 2020; McKee et al., 1986).

**Figure 3.**
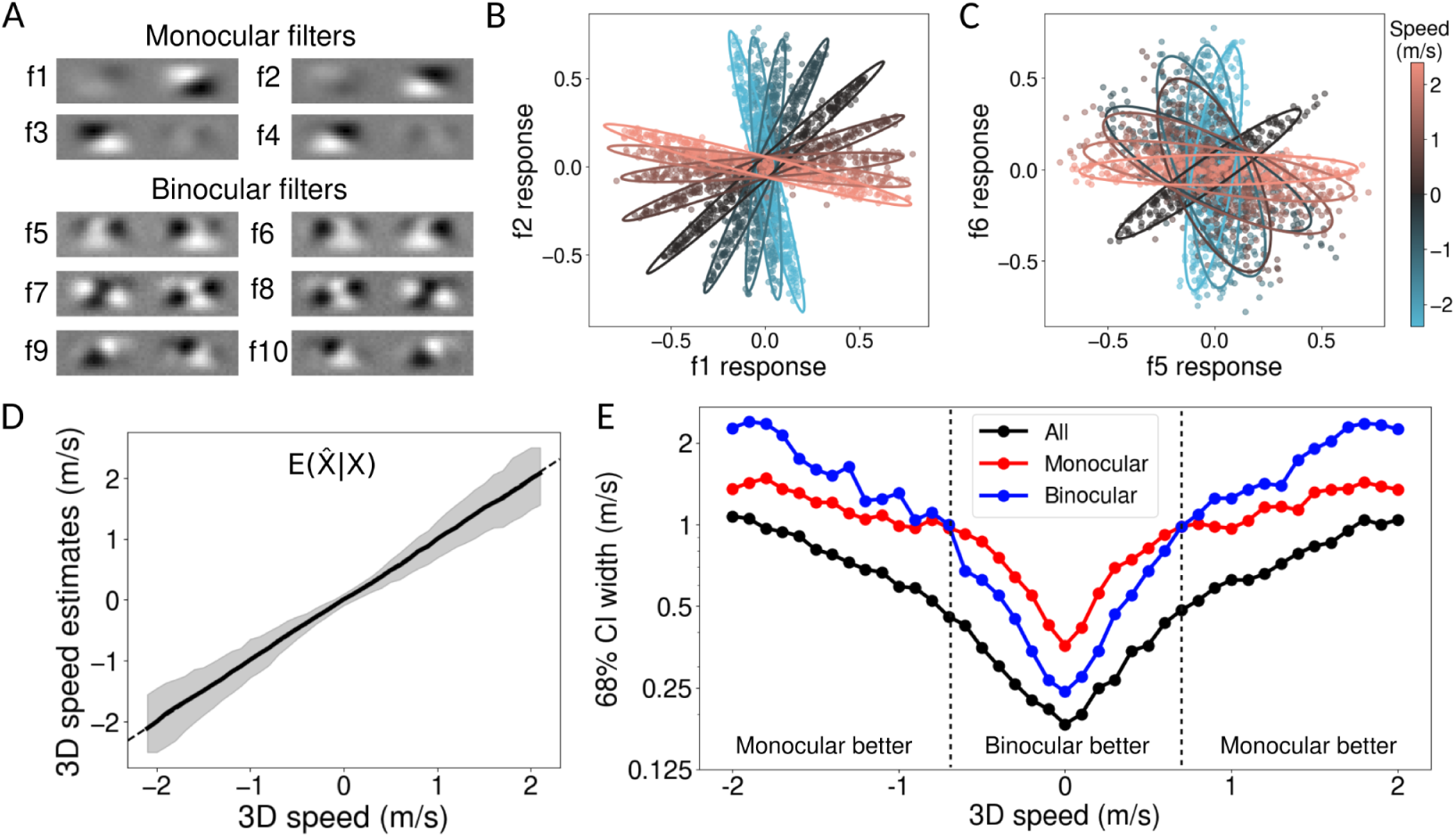
Task-optimized filters, responses, and 3D speed estimation performance**. A,** Optimal binocular spatiotemporal filters for 3D speed estimation. Two specialized filter populations spontaneously emerged. Monocular filters (top) support the extraction of inter-ocular-velocity-difference (IOVD) cues (see main text). Binocular filters (bottom) support the extraction of changing-disparity-over-time (CDOT) cues (also see main text). **B,** Conditional filter responses for the first pair of monocular filters in A for a subset of the 3D speeds in the dataset (colors). Each symbol is the expected filter response to an individual stimulus (i.e. without added filter noise). Ellipses show the best-fitting Gaussians. **C,** Same as B but for a pair of binocular filters. **D,** Median estimated 3D speeds as a function of true 3D speed. Grey area shows 68% confidence intervals of the response distribution. **E,** Confidence intervals on estimates derived from monocular filters alone (red), binocular filters alone (blue), or all filters together (black). Dashed lines indicate speed ranges where monocular filters and binocular filters are better.

The filters that optimize 3D speed estimation performance have some interesting properties. Two distinct filter-types were spontaneously learned for the task (**Figure 3A**). The first four filters are monocular; they assign strong weights to one eye and weights near zero to the other eye (**Figure 3A top**). The fact that monocular filters are learned may be surprising given that the inputs to all filters were binocular videos. (The emergence of monocular filters is non-trivial. Performing PCA on the dataset, for example, does not return monocular filters.) The remaining six filters are binocular; they weight information from each of the two eyes approximately equally, with left- and right-eye components that select for the same spatial frequency (**Figure 3A bottom**). The variation in ocular dominance across the filters may well have functional importance for this task (see below). Variation in ocular dominance is a well known property of cortical neurons (Hubel & Wiesel, 1962; Katz & Crowley, 2002). Also, the binocular receptive fields have the interesting property that the position of each eye’s preferred feature changes smoothly in opposite directions across time, entailing that the preferred binocular disparity changes across time. This property is the signature of receptive fields that jointly encode motion and disparity. The presence of such receptive fields in early visual cortex, and their potential functional importance has been written about extensively (see Discussion).

To understand why two filter types are learned, we examined how each type supports 3D speed estimation. We examined estimation performance for versions of the model that used only the monocular filters, or only the binocular filters, and compared the precision of the estimates obtained with each model across speeds. At slow 3D speeds, estimates derived from the six binocular filters alone have smaller confidence intervals (i.e. are more precise) than the estimates derived from the four monocular filters alone. At fast speeds, the opposite is true (**Figure 3E**). (Similar results are obtained when the number of filters are matched between the two groups–for example, four binocular filters vs. four monocular filters). When all filters are used together, the confidence intervals are more similar to those of the monocular filters at fast speeds, and are more similar to those of the binocular filters at slow speeds. Each filter type has a domain of specialization. Thus, the two filter types are engaged in a division of labor. The filters extract complementary task-relevant information from the stimuli.

Interestingly, the two distinct filter types appear to support extraction of two different 3D motion cues used by humans and non-human primates in 3D motion estimation: the interocular velocity difference (IOVD) cue and the changing disparity over time (CDOT) cue (Cormack et al., 2017). The IOVD cue is the difference in the speeds of the retinal images of the moving object in the two eyes, produced by motion in depth. Computing this cue involves estimating the velocity of each retinal image first, and then determining the velocity differences between the eyes. The monocular filters are well-suited to this computation. The CDOT cue is the change in binocular disparity over time as an object moves in depth. Computing this cue involves computing the binocular disparity at each time point, and then determining how the disparity changes with time. The binocular filters are well-suited to this computation.

Although in natural environments moving objects tend to produce IOVD and CDOT cues that are consistent with one another, processing of the two cues requires different neural computations. Psychophysical and neurophysiological investigations have provided strong evidence that IOVD and CDOT cues are both used by the visual system (Cormack et al., 2017). Specialized IOVD- and CDOT-isolating stimuli have been used to show that the two cues underlie different 3D motion sensitivity profiles under different stimulus conditions. Humans rely on IOVD cues more heavily at high speeds and in the peripheral visual field, and on CDOT cues more heavily at low speeds and in the fovea (Cormack et al., 2017; Czuba et al., 2010a). It is interesting that ideal observer performance is supported by the distinct filter types in a manner that dovetails with human psychophysical results. These similarities between human and ideal observer 3D speed estimation performance suggest that the adaptive manner in which humans use these motion cues across conditions may reflect near-optimal processing of the available visual information.

There are additional points to make about how the filters respond to naturalistic stereo movies. First, the information about 3D speed is carried near-exclusively by the covariance of the filter responses, because the mean filter responses are essentially equal to zero for all speeds. Second, the filter responses, conditioned on each 3D speed, are tightly approximated by Gaussian distributions (**Figure 3B,C**). These results are important on methodological grounds because they justify the use of AMA-Gauss, which approximates each conditional response distribution *p*(*R|X_j_* ) as Gaussian during the filter learning process (Jaini & Burge, 2017). More fundamentally, these results indicate that quadratic combination of the filter responses is required for computing the likelihood of the different speeds, L(X_i_|**R**), and thus for optimal inference of local 3D speed. Taking the logarithm of the expression for the likelihood (Equation 4), shows that the log-likelihood of a given 3D speed is given by a quadratic combination of the filter responses:

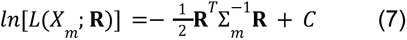

(for clarity we omit the response means _µ*m*_ from the equation, since they are nearly zero for all speeds). These optimal computations are closely related to those posited by the energy model, a popular descriptive model of complex cells.

The energy model gets its name because it computes the local energy (phase-invariant content) in different spatiotemporal frequency bands (Adelson & Bergen, 1985). It achieves this phase-invariance by first computing the output of a pair of orthogonal quadrature filters, and then squaring and adding together these outputs. Energy-like computations underlie models of neural response to arbitrary contrast patterns (Park et al., 2013; Rust et al., 2005), and models of neural selectivity for motion-in-depth (Wu et al., 2020), 2D motion (Adelson & Bergen, 1985), and disparity (Burge & Geisler, 2014; DeAngelis et al., 1991; Ohzawa, 1998). Such computations have also been shown, with natural stimuli, to support the optimal estimation of many of these same latent variables, including 2D motion (Burge & Geisler, 2015; Chin & Burge, 2020), binocular disparity (Burge & Geisler, 2014; Jaini & Burge, 2017), and focus error (Burge & Geisler, 2011, 2012). Here, from a task-specific analysis of natural signals, we have shown that they support the optimal estimation of 3D speed.

Unlike the energy model, however, the filters in our observer model are not constrained to be a quadrature pair. Because of how feature selectivity interacts with the effects of internal noise, filter pairs that span the same subspace as a quadrature pair but are non-orthogonal can, under certain circumstances, produce a higher quality encoding than orthogonal filter pairs (Burge & Jaini, 2017). Also, the quadratic computations that are optimal for probabilistic decoding are performed with a specific set of weights given by the inverse covariance matrix Σ_m_^-1^, which are, in turn, determined by natural scene statistics (**Figure 1B**). Thus, the optimal computations in our model can be implemented by likelihood-neurons that are a generalization of the energy model of complex cells. The current results therefore provide a normative explanation, grounded in natural scene statistics, for the success of a common descriptive model of neural response. These results are in line with similar results found for other simple tasks (Burge, 2020).

### 3D-direction estimation

Next, we developed an ideal observer for the estimation of 3D-direction from naturalistic binocular videos. The results for the 3D-direction estimation task are similar in many ways to the 3D-speed estimation results. The filters that optimize 3D direction estimation, their responses to naturalistic stimuli, and the ideal observer estimates of 3D direction are shown in **Figure 4**.

**Figure 4.**
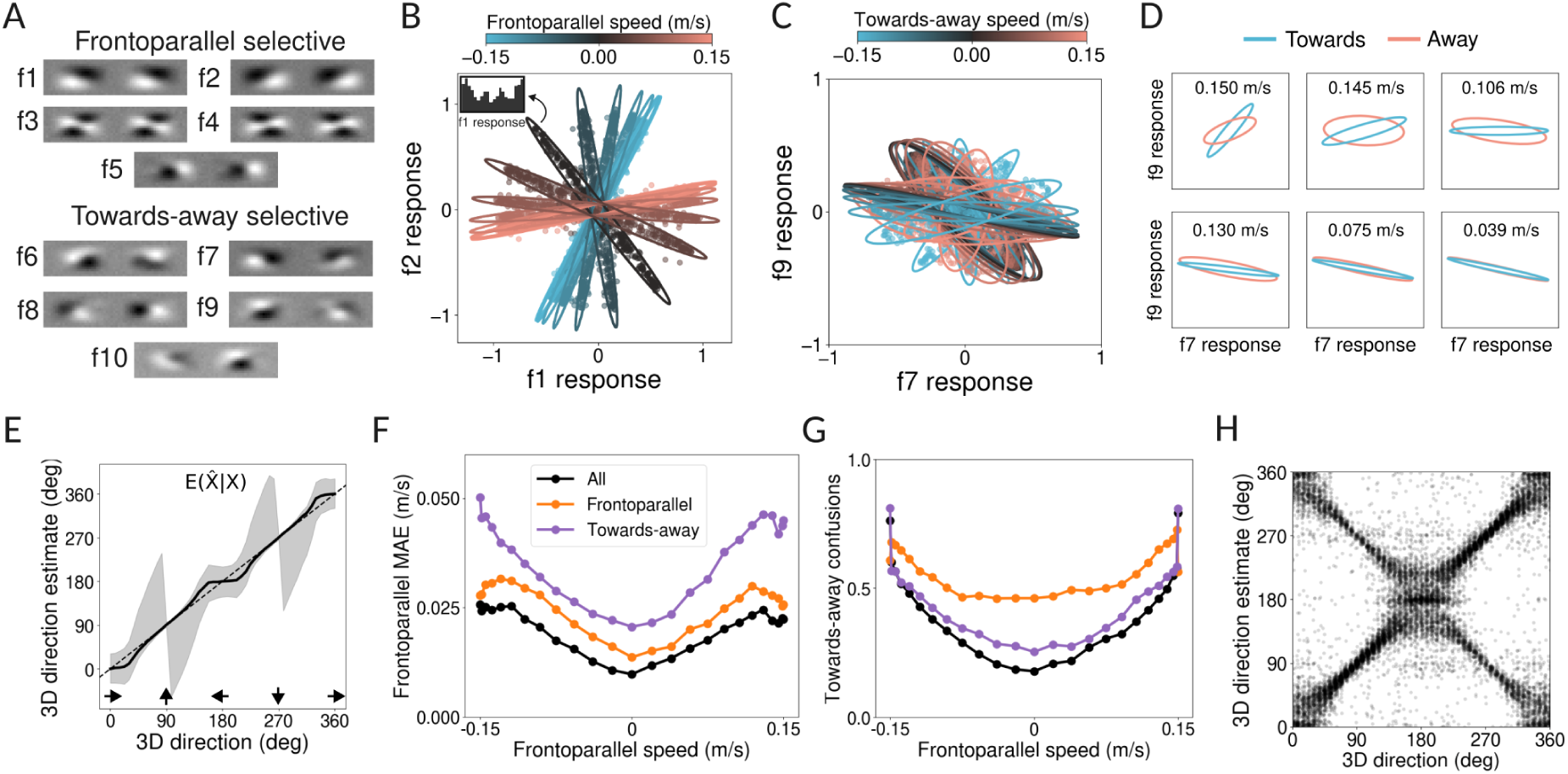
3D direction estimation **A,** Filters learned for 3D direction estimation. Top: Frontoparallel selective. Bottom: Towards-away selective. **B,** Response distributions for the pair of frontoparallel selective filters, conditional on 3D direction, and color coded by the frontoparallel motion component (only a subset of the directions). Ellipses show the best-fitting Gaussians. Filter responses without added noise are shown. 3D directions with identical frontoparallel components but opposite towards-away components produce perfectly overlapping response distributions for this pair of filters. Inset shows a response histogram for filter **f**_1_, for the 90° direction. **C,** Same as B, but for towards-away filters, color coded by their towards-away motion component. **D,** Same as C, each panel shows only a direction pair that have the same frontoparallel component but opposite towards-away component. Panel headers indicate the magnitude of the towards-away speed for each pair. The color of the ellipse indicates whether the frontoparallel component is towards or away from the observer. **E,** Estimated 3D directions. Gray area shows 68% confidence intervals. **F,** Proportion of towards-away sign confusions obtained from the different types of filters. Confusion rates of over 0.5 are possible for some motions because the sign could be towards, away, or 0 (e.g. frontoparallel motion). **G,** Mean absolute error of the frontoparallel component of estimated motion as a function of true 3D motion direction. **H,** Model estimates across the dataset. Each point shows the estimate from a unique stimulus.

Two filter sub-populations are again learned for the task. First, some filters have almost identical patterns of weights in the two eyes (**Figure 4A top**), while others have quite distinct patterns of weights (**Figure 4A bottom**). Second, we observed that no pair of filters clearly discriminate the full range of 3D directions Rather, different filter pairs specialize in discriminating different cardinal directions. Filtering having the same weights in the two eyes yield conditional response distributions that are segregate 3D directions with different frontoparallel frontoparallel component (**Figure 4B**). But, for a fixed frontoparallel speed, these filters yield response distributions that perfectly overlap for different towards and away directions (**Figure 4B**). The filters with distinct weights in the two eyes do not clearly segregate directions with different frontoparallel component, but they do discriminate motions towards and away from the observer (**Figure 4C**). In particular, when conditioned on the true frontoparallel component, responses clearly segregate according to the towards and away directions, and the segregation decreases with the magnitude of the towards-away component (**Figure 4D**). The results above are consistent with previous work showing that binocular neurons with identical selectivities in the two eyes cannot discriminate towards-away direction with non-looming stimuli such as the ones used here (Bonnen et al., 2020). Thus, we call these filter populations frontoparallel and towards-away filters respectively.

In line with the observations above, when estimating 3D direction with frontoparallel filters, the frontoparallel motion component of the estimate is accurate, but this is not the case for the towards-away filters (**Figure 4F**). On the other hand, direction estimates derived exclusively from the frontoparallel filters confuse the sign of towards-away motion approximately 50% of the time (**Figure 4G**, orange). Estimates derived from the towards-away filters confuse the sign a much smaller percentage of the time (**Figure 4G**, purple), but have a high error. The two types of filters have clear functional specializations.

Interestingly, although one may expect the towards-away filters learned for this task (**Figure 4A bottom**) to be similar to the 3D speed filters that discriminate different speeds in the sagittal plane (**Figure 3A**), the two populations are different. However, this is not surprising if we consider the many differences between the two tasks (e.g. the range of towards-away speeds, the added frontoparallel components in the direction task) and the different functions of the two filters (estimate 3D speed and discriminate towards and away motion).

Again, like with 3D speed estimation, the response distributions are reasonably approximated as Gaussian and nearly all of the task-relevant information is carried by the covariance of the filter responses (**Figure 4BC**). However, some filter responses are clearly non-Gaussian (**Figure 4B inset**). This finding suggests that more sophisticated computations than the energy-model-like computations described here are required to extract all available information. However, because the Gaussian approximation is reasonably good and captures all task-relevant information up to second-order (i.e. the covariance of filter responses), the performance differences between quadratic decoding and optimal decoding will be minor.

Finally, again like in the 3D-speed estimation task, ideal observer performance in the 3D-direction estimation task is similar to key aspects of human psychophysical performance. For a non-negligible proportion of the stimuli, the ideal observer confuses towards and away motion directions (**Figure 4H**), generating a characteristic X-shaped pattern of responses. Though counter-intuitive, this surprising pattern of responses is characteristic of human performance in laboratory based 3D motion tasks (Bonnen et al., 2020; Fulvio et al., 2015; Rokers et al., 2018), indicating that this type of error may be a consequence of optimal decoding of 3D direction from stereo-based cues to 3D motion. Furthermore, for some directions, model estimates are biased towards frontoparallel directions (**Figure 4EH**, note the response cluster around 180 deg). Frontoparallel bias is a known feature of human 3D direction discrimination that has been previously attributed to slowness priors using ideal observer analysis (Rokers et al., 2018; Welchman et al., 2008). Here, however, we show that the same behavior can be obtained in an ideal observer that does not include a slowness prior (also see Rideaux & Welchman, 2020).

### Disparity variability affects optimal 3D motion processing

We next analyzed the effect of adding an additional source of nuisance variability to the task: disparity variability. In the previous sections all videos started from zero disparity, which is appropriate for perfectly fixated and foveated objects. However, small convergent or divergent eye movements cause fixation errors that are known as vergence noise, or vergence jitter. Under typical conditions, the standard deviation of vergence noise has been estimated to range between 2 arcmin and 10 arcmin, and is known to harm stereo-based depth discrimination performance (Ukwade et al., 2003b, 2003a). We examined the effect of disparity variability across, and exceeding, this range on 3D motion estimation performance. (Note that decoding routines used response statistics appropriate to the modified dataset.)

For 3D speed estimation, disparity variability reduces estimation precision for low speeds (**Figure 5A**) but not for fast speeds (**Figure 5B**). The reduction in precision at low speeds occurs because disparity variability strongly affects the information provided by the binocular filters; it has little influence on the information provided by the monocular filters (see **Figure 5A**). At high enough levels, disparity variability eliminates the advantage of binocular filters over monocular filters at low speeds. For 3D direction estimation, disparity variability increases towards-away confusions (**Figure 5C**). It has only a small effect on the frontoparallel component of the direction estimates (**Figure 5D**).

**Figure 5.**
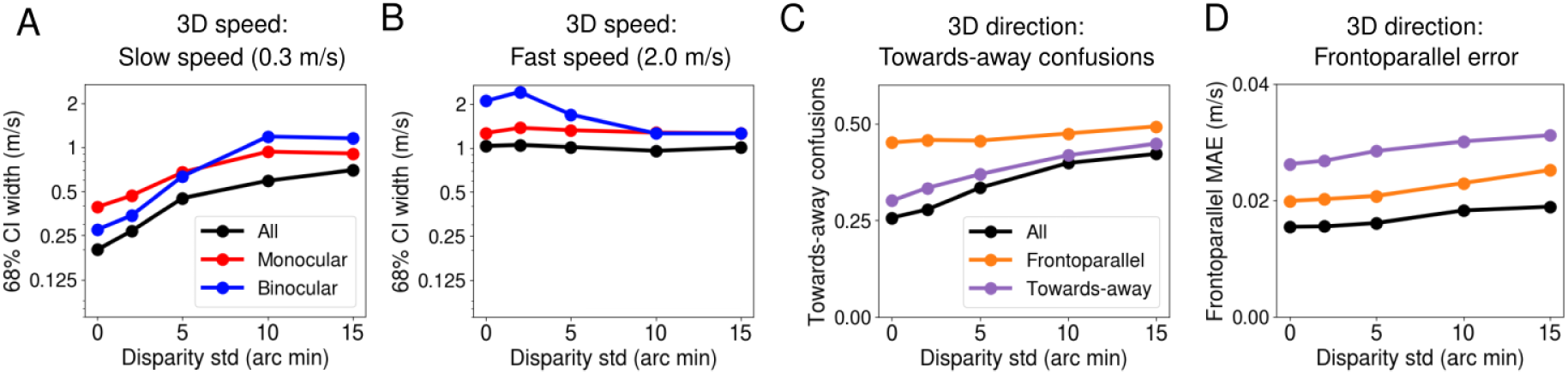
Disparity variability effects are condition-specific. **A,** Confidence intervals of the 3D speed estimates, as a function of disparity variability for a low target speed (0.3 m/s). The filter subset used to obtain the estimates is indicated by line color. **B,** Same as A for a fast target speed (2.0 m/s). **C,** Proportion of towards-away sign confusions as a function of disparity variability, for a target direction of 22.5° and speed of 0.15 m/s. The filter subset used to obtain the estimates is indicated by line color. **D,** Mean absolute error of the estimated frontoparallel motion component as a function of disparity variability, for a target direction of 22.5°.

There are at least two ways in which these effects may be relevant for understanding human psychophysical patterns of performance. First, it is well established that, for 3D towards vs. away motion perception, peripheral vision relies more strongly on IOVD cues than the fovea (Cormack et al., 2017; Czuba et al., 2010a). And it has been shown that, in natural viewing, the peripheral visual field has higher disparity variability than the fovea (Sprague et al., 2015). The results in **Figure 5** thus suggest that the periphery may preferentially rely on IOVD cues in part because of higher disparity variability. However, because there are other relevant differences between the foveal and peripheral visual fields–for example, fast retinal motion speeds are more common in the periphery than at the fovea (Matthis et al., 2022)–a dedicated modeling effort would have to be undertaken to understand their relative importance. Still, the **Figure 5** results are intriguing, and may help explain functional differences between foveal and peripheral vision. Second, a study on 3D direction estimation showed that when initial target disparity is different from zero, towards-away confusions substantially increased, while left-right confusions were not affected (Fulvio et al., 2015), similar to our results.

## Discussion

Despite the importance of 3D motion estimation and discrimination for behavior, relatively little is known about the computations that the brain uses, or what computations are optimal, for estimating 3D motion from the retinal input. To gain insight into the normative computations that the brain should use, we developed image-computable ideal observers grounded in the task-relevant natural scene statistics for each of two tasks: 3D speed estimation and 3D direction estimation. For both tasks, two populations of filters spontaneously emerge from a Bayesian filter-learning routine, each with its own functional specialization. For 3D speed estimation, the filter populations each relate to one of two well-characterized binocular cues to 3D motion (IOVD and CDOT cues), providing a normative explanation for their use by humans. For 3D direction estimation, one filter population strongly selects for image-features supporting estimation of left-right motion components; another provides useful information for discriminating towards and away motion. No such populations are described in the literature. The present work motivates future neurophysiological experiments. We also found that a generalization of the energy model of complex cells could implement optimal inference for the 3D speed estimation task, and closely approximate optimal inference for 3D direction estimation. Finally, ideal observer estimation performance in each task mirrors intriguing–and sometimes counterintuitive–patterns of human performance, suggesting that human performance patterns may be accounted for by the ideal computations.

### Limitations and future directions

The tasks upon which the analyses of 3D motion estimation were based are limited in some respects. The binocular retinal videos that the ideal observers used to estimate 3D motion depicted surfaces that started moving 1m away, straight-ahead (see **Figure 2B**). To obtain an unambiguous estimate of 3D velocity from a pair of left- and right-eye retinal image motions, information about the 3D location (i.e. distance and direction) of the target is required. Any given binocular retinal video is compatible with many different 3D motions, depending on the starting location of the target (Lages & Heron, 2010). Under certain cue-impoverished circumstances, in which location information is poor, humans cannot accurately estimate 3D motion (Rushton & Duke, 2009). Under typical conditions, however, location information can be extracted from a combination of image-based and extra-retinal cues (Backus et al., 1999; Watt et al., 2005). When 3D location is accurately estimated, the accuracy of stereo-3D-motion estimates are limited by the accuracy with which binocular retinal motion is estimated.

The current efforts to model 3D motion estimation do not address how 3D location is determined by the visual system. But, just as the core computation underlying stereo-based depth estimation–solving the stereo-correspondence problem (Burge & Geisler, 2014; Read & Cumming, 2007; Tyler & Julesz, 1978)– does not depend on location estimation, the core computations described here do not depend on location estimation. Hence, the computations described here provide the optimal solution to a critical sub-task required for estimating motion-in-depth, just as solving the correspondence problem is a critical sub-task for disparity-based depth estimation (also see Methods: Caveats).

Another limitation is that we analyzed 3D speed estimation and 3D direction estimation separately. That is, similar to many psychophysical and neurophysiological studies, we examined estimation of multiple speeds for a fixed direction, and estimation of multiple directions for a fixed speed. However, it is possible that estimating direction and speed simultaneously requires different optimal features or strategies than estimating them separately. For example, the task of estimating 3D speed for a fixed direction can be solved by estimating monocular 2D speed, because at a given 3D location there is a one-to-one mapping from 3D speed to retinal speeds (also see Methods: Caveats). When the task requires the joint estimation of 3D speed and direction, this particular issue is no longer present. Future work will examine 3D motion estimation as a single task in which both speed and direction can vary. Because of the strong tradition of studying 3D speed and 3D direction separately, there is obvious value in understanding how the optimal solutions to the isolated tasks are related to the optimal solution 3D velocity estimation tasks that characterize everyday viewing situations (Burge et al., 2019).

Further, the binocular videos in the main stimulus set lacked some 3D motion cues that are present in natural stimuli, chiefly looming cues. Looming cues are changes in the retinally-projected sizes of environmental features as they change distance from the observer. We generated stimuli that include looming cues, and found that all patterns described above–filters, response distributions, estimation performance–are reproduced with these stimuli, except that estimation performance was marginally more precise. However, because all stimuli contained only vertically-oriented stimulus information, the usefulness of looming cues are probably underestimated here. Future work should analyze binocular videos without vertical averaging, so all orientations are available, to better understand the relative contributions of the different cues available in more realistic stimuli.

### Joint encoding of motion and disparity

The neurophysiological underpinnings of 3D motion perception is a topic of recurring interest (Rosenberg et al., 2023). The results here indicate that binocular receptive fields that jointly encode retinal speed and disparity information optimally support the estimation of 3D speed (**Figure 3A**, bottom), especially at slow speeds (see **Figure 3E**). Models based on complex cells having subunit receptive fields that jointly encode motion and disparity have been proposed to account for neural activity underlying 3D motion estimation performance (Anzai et al., 1999; Pack et al., 2003; Qian & Andersen, 1997). However, direct neurophysiological measurements have produced scant evidence for neurons that jointly select for motion and disparity, or that explicitly code for 3D motion, in early visual cortex (Read & Cumming, 2005). Similarly, evidence that such neurons exist in area MT is not iron-clad (Czuba et al., 2010b; Sanada & DeAngelis, 2014; Thompson et al., 2023). The paucity of these neurons in these cortical areas, of course, does not mean that humans cannot estimate motion-in-depth. Clearly, humans can estimate 3D motion. And neurons that unambiguously select for 3D motion have recently been reported in area FST (Rosenberg et al., 2023; Thompson et al., 2023).

There are loosely analogous results in stereo-surface-orientation perception. There is no strong evidence that, in the early visual system, there exist binocular receptive fields that are optimized for estimating 3D surface orientations different from frontoparallel, even though such receptive fields have been searched for specifically (Banks et al., 2004; Bridge et al., 2001; Bridge & Cumming, 2001; Greenwald & Knill, 2009; Nienborg et al., 2004). Of course, whether or not such binocular receptive fields exist in early visual cortex is not determinative of whether or not humans can estimate 3D surface orientation (i.e. the slant and tilt). Humans can clearly do so (Hillis et al., 2004; Kim & Burge, 2018, 2020; Knill, 1998; Stevens, 1983; Watt et al., 2005), and there are cortical areas that contain neurons that jointly select for slant and tilt (Rosenberg et al., 2013; Rosenberg & Angelaki, 2014).

So, what is the utility of this discussion, and what lessons can be taken from the present ideal observer analysis? In 3D surface orientation estimation, the lack of early optimal detectors entails that the rest of the visual system will be subject to the limits imposed by those non-optimal early detectors (see Banks et al., 2004 for a nice discussion of this issue). Similarly, the apparent rarity of neurons with joint motion-disparity selectivity in the early primate visual system suggests that humans may be subject to deficits in 3D motion perception that the ideal observer may not be. Targeted experiments could be developed to test whether those deficits in fact occur. Such experiments should be conducted at slow 3D speeds, because it is at these speeds that the binocular receptive fields place limits on performance.

### Priors and perceptual estimation

We have developed ideal observer models for the estimation of 3D motion. These models are grounded in natural image statistics, incorporate front-end properties of the visual system, and make use of the tools of probabilistic inference. Unlike many probabilistic models of motion perception (Rokers et al., 2018; Stocker & Simoncelli, 2006; Weiss et al., 2002; Welchman et al., 2008), however, we did not impose a zero-motion prior. We imposed a flat prior instead, consistent with the relative number of stimuli at each level of the latent variable in the stimulus set. Estimation errors nevertheless emerged that are typically attributed to the action of a zero-motion prior. Two examples are as follows. First, estimates of 3D direction were biased towards the fronto-parallel direction. Second, estimates of 3D speed from low-contrast stimuli in the dataset tend to be more biased towards slow speeds (results not shown). Both the first finding, and an analog of the second finding in 2D motion estimation, have been appealed to as evidence for the action of a slow-motion prior in perceptual estimation. The current results indicate that the same patterns of estimation errors can emerge from an accuracy-maximizing ideal observer with a flat prior. The current results suggest that patterns of estimation errors that have been attributed to priors can also be caused by properties of natural stimuli and/or constraints imposed by the front-end of visual systems that have nothing to do with priors. The current findings thus invite reevaluation of common claims in the literature that perceptual biases constitute evidence for non-flat priors (also see Rideaux & Welchman, 2020).

## Conclusion

A spate of recent research has investigated the psychophysical limits and neurophysiological underpinnings of 3D motion estimation (Cormack et al., 2017; Rosenberg et al., 2023). The development of ideal observers for 3D motion estimation–that are grounded in natural scene statistics–can help interpret existing results by providing principled benchmarks against which performance patterns can be evaluated. They can also suggest new hypotheses that can be tested experimentally, or provide reasons to question consensus views about how to best understand widely observed neural and perceptual phenomena (Burge, 2020). The ideal observers developed here, for the tasks of 3D speed and 3D direction, show that many, sometimes confusing, aspects of neural activity and human performance may be a consequence of optimal information processing in the visual system. Increasing the realism of the stimulus sets and the generality of tasks should provide deeper insights still into the computations and performance patterns that characterize 3D motion perception in natural viewing.

## Acknowledgements

This work was supported by the National Eye Institute and the Office of Behavioral and Social Sciences Research, National Institutes of Health Grant R01-EY028571 to J.B.

## Author contributions

D.H and J.B designed the research, D.H wrote the analysis code, D.H and J.B analyzed the data, D.H and J.B wrote the first draft of the paper, D.H and J.B edited the paper.

